# Pangebin: improving plasmid binning in bacterial isolates using pangenome-assembly graphs

**DOI:** 10.1101/2025.04.10.648291

**Authors:** Mattia Sgro, Brona Bejova, Yuri Pirola, Tomas Vinar, Paola Bonizzoni, Cedric Chauve

**Affiliations:** Department of Informatics, Sistems and Communication, University of Milano - Bicocca, Milan, Italy; Faculty of Mathematics, Physics and Informatics, Comenius University in Bratislava, Slovakia; Department of Mathematics, Simon Fraser University, Burnaby, BC, Canada

## Abstract

Short-read genome assemblies typically consist of many contigs of variable lengths and their putative connections represented as an assembly graph. Assembly graphs produced by different tools from the same data may differ significantly, posing a challenge to tools for downstream processing tasks. One such task is plasmid binning, that is identifying plasmids in sequenced bacterial isolates, which is crucial for monitoring the spread of antimicrobial resistance. When plasmid binning tools are applied to assembly graphs produced by different tools, they may exhibit different performance, and choosing the best results a priori can be difficult. To address the above issue, we propose the use of a pangenome graph, built from assembly graphs produced by assembling short reads of the same sample with different assemblers. The resulting pangenome-assembly graph highlights similarities between contigs from different assemblies while retaining information on contigs that appear only in one of the input assemblies. We then used the `PlasBin-flow` plasmid binning tool customized to take into account pangenome information to identify plasmid bins. The results for pangenome-assemblies built by Unicycler and Skesa show an increase in accuracy measures compared to the mean results obtained on single assemblies, leading to an overall more accurate prediction than a blind choice of assemblers. The source code of the pipeline is available at https://github.com/AlgoLab/pangebin along with the dataset used in this study.

## 1 Introduction

Plasmids contribute to the propagation of genes responsible for antimicrobial resistance (AMR) in bacteria [1]. Due to the emergent relevance of AMR in public health surveillance [2, 3], the development of computational methods that are able to detect plasmids from sequencing of bacterial genomes is of great interest. Short-read sequencing is the most common low-cost technology used for microbiome genomics surveillance. In this framework, short reads from a bacterial isolate are assembled into contigs to face three computational problems in plasmid identification: classification, binning, and assembly. In particular, the classification problem aims to distinguish plasmid and chromosomal contigs, the binning problem aims to cluster together plasmid contigs into bins, each representing a specific plasmid, and the assembly problem aims to reconstruct the whole plasmid sequence. In this work we focus on the plasmid binning problem. Current state-of-the-art tools are aimed at extracting plasmids from assembled contigs of a sequenced bacterial genome are based on de novo methods, reference-based strategies, or a combination of both. While reference-based methods, such as MOB-recon [4], make use of homology to databases of known plasmids, de novo approaches, such as Recycler [5], PlasmidSPAdes [6] and gplas [7, 8, 9] strongly rely on contig features such as length, coverage or K-mer count, circularity and GC-content. Moreover, gplas also leverages machine learning approaches to assign a plasmid score to a contig, i.e., the probability of that contig to be a plasmid, via mlplasmids [10]. Hybrid methods like HyAsP [1], Plasbin [11] and `PlasBin-flow` [12] combine the use of a reference database of known genes with the exploitation of contig features described above. In general, plasmid binning tools rely on analyzing the assembly graph of a bacterial sample and these graphs are generally obtained by so-called assemblers, such as Unicycler [13], Skesa [14], or PlasmidSpades [6]. Note that some binning tools are tailored to work on graphs built by specific assembler(s); for example `gplas` accepts exclusively Unicycler graphs, whereas there are assembler-agnostic tools, such as `PlasBin-flow`, that work on any kind of assembly graph. In this work we tested these two binning tools along with Skesa and Unicycler assembly graphs. Note that in order to test `gplas` with Skesa assemblies, we had to convert assembly graphs obtained with Skesa into Unicycler graph format. Comparing these two tools we observed that `PlasBin-flow` performed generally better with Skesa graphs but obtained better recall with Unicycler, while `gplas` showed the opposite, performing better with Unicycler but obtaining better precision with Skesa. Moreover, there were also samples that did not show this mean behavior, and we concluded that -a priori-it is not possible to choose in absolute means the best performing assembler for a particular binning tool.

In this paper, we address this limitation by proposing the construction of a pangenome-assembly graph that combines contigs from different assembly graphs built from the same bacterial isolate. More precisely, our goal is to have a comprehensive representation of information contained in multiple graphs, showing overlaps and links between contigs of different assembly graphs. By leveraging the recent advances in multiple genome sequence comparisons based on the notion of a pangenome, in this work, we propose the idea of building a *pangenome-assembly graph*. Observe that the notion of a pangenome graph is traditionally referred to the representation of a collection of multiple genomes [15, 16] that are either compared through an all-versus-all alignment or aligned over a reference sequence. In this paper we extend this view, considering the pangenome built from a collection of sequences of a single sample, extracted by multiple assembly graphs built using different assembly tools. This novel representation incorporates the original edges of the input assembly graphs into a pangenome graph built through an all-versus-all alignment of the contigs of two or more assembly graphs. It may also include information necessary for downstream analysis (in our case, plasmid binning), which was also inherited from the original assembly graphs. Furthermore we propose an adaptation of `PlasBin-flow` to work on pangenome-assembly graphs to provide a novel approach to plasmid binning that works from assembly graphs coming from different assemblers: our strategy is implemented in the tool `Pangebin`. Observe that `Pangebin` method exploits the pangenome capabilities to aggregate information from each assembly graph, providing a solution that improves over the best solution that could be achieved by analyzing each assembly graph individually.

To build our pangenome graph we relied on `nf-core/pangenome` [17], a well-established pipeline for building pangenome graphs based on `PGGB` [16]. Starting from a set of contigs coming from multiple assembly graphs, it returns a pangenome with contigs split into overlapping nodes and connected by paths in the graph. In this work we combine assembly graphs built with Skesa [14] and Unicycler [13]. The graph built from `nf-core/pangenome` was then augmented with information from the input assembly graphs such as original edges linking contigs and contig features relevant for the binning task, such as GC-content, gene density, coverage, and circularity.

We tested `Pangebin` on several bacterial samples and observed that overall accuracy improved over the mean of the accuracy obtained by `PlasBin-flow` on each assembly. In particular our tool performs better in precision than `PlasBin-flow` run on a Unicycler assembly, while also performing better in recall than `PlasBin-flow` run on Skesa assembly graphs. Moreover we claim that in a scenario in which we do not know a priori which assembly graph provides the best performances for a given sample, our approach is indeed more robust than a blind choice of the assembler.

## 2 Method

The novelty of our plasmid binning tool is that it leverages the concept of pangenome graph to build a consensus graph (that we call *pangenome-assembly graph*) starting from different assembly graphs. Although `Pangebin` can be potentially used for any assembly graph, in this work, the input data for `Pangebin` is given by the assembly graphs produced by the two most common assemblers used in plasmid analysis: Skesa [14] and Unicycler [13]. Then, any plasmid binning tool can be applied to the pangenome-assembly graph provided it does not rely on specific assumptions that may not hold for such graphs. In this work, we adopted `PlasBin-flow` [12] as the plasmid binning tool since `PlasBin-flow` solves the binning problem using an MILP formulation that is particularly well-suited to integrate additional properties that stem from pangenome-assembly graphs. Indeed, we extended the MILP formulation to incorporate the preference of selecting nodes (contigs) in the plasmid binning that are supported by both assembly graphs, thereby likely improving its accuracy.

The methodology implemented in `Pangebin` consists of the following three steps:

1. a pair of assembly graphs is used to build a pangenome graph representing the similarities among the input contigs;
2. the pangenome graph is transformed into a pangenome-assembly graph by enriching it with information from the original assembly graphs;
3. a tailored version of `PlasBin-flow` is applied to the pangenome-assembly graph to obtain plasmid bins.

### 2.1 The pangenome graph *G*_*p*_: a consensus of sequence contigs

The input to our method consists of a set of assembly graphs built from the same set of short reads. For simplicity, we illustrate the method considering only two generic assemblers, *a* and *b*, but the method can be easily extended to any set of short-read assemblers. Suppose that the two assemblers respectively build the two assembly graphs *G*_*a*_ = (*C*_*a*_, *E*_*a*_) and *G*_*b*_ = (*C*_*b*_, *E*_*b*_), with *C*_*a*_ (*C*_*b*_, respectively) being the set of contigs of graph *G*_*a*_ (*G*_*b*_, respectively) and *E*_*a*_ (*E*_*b*_, respectively) the set of edges linking contigs in the assembly graph. An edge connecting two contigs *c*^*′*^ and *c*^*′′*^ is represented as a pair 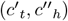, meaning that we are linking the *tail* of the first contig with the *head* of the second contig.

The first step is to build a pangenome graph *G*_*p*_ from the whole set of contigs *C* = *C*_*a*_ ∪ *C*_*b*_ extracted from the assembly graphs *G*_*a*_ and *G*_*b*_ using the *nf-core/pangenome* pipeline [17], which is based on the PanGenome Graph Builder tool (PGGB) [16]. Observe that graph *G*_*p*_ is the result of a reference-free pairwise comparison of the contig set, as we detail below. Given a set of input sequences, PGGB performs an all-versus-all sequence comparison of the sequences. This comparison aims at identifying substrings of the input contigs that have significant similarity, namely at least 95% base-pair similarity. The above percentage is the one suggested by the default parameters of PGGB tool. Moreover, in order to avoid matching small contigs together, we set a threshold of 200bp as the minimum length to perform the matching. This choice is due to the fact that otherwise PGGB tool creates bubbles for SNP (Single Nuleotide Polymorphisms) that are ignored by our approach. Contigs that are similar according to these criteria are then split into substrings called *fragments*. Contigs that are similar according to these criteria are then split into *fragments*. In particular, when two or more contigs share a common subsequence, we split those contigs in at least one *shared* fragment and eventually some *private* fragments. In particular shared fragments contain the sub-sequences common to all the matching contigs and are linked to each non-matching sequence of the similar contigs, represented by private fragments. Note that, in case of perfect matching of two or more contigs, a single shared fragment is produced and no private fragment is produced; in general, higher the similarity between two contigs, less is the length of the produced private fragments. Finally, each fragment is associated with a distinct node of the graph. Hence, the set *F* of fragments forms the set of nodes of the pangenome graph *G*_*p*_ = (*F, E*_*p*_) while the set *E*_*p*_ contains edges that connect nodes based on the contiguity of fragments in the input contigs. More precisely, there exists an edge between two nodes 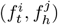 (tail and head of fragments *f*^*i*^ and *f*^*j*^, respectively) if *f*^*i*^ and *f*^*j*^ are consecutive fragments of some contig *c* ∈ *C*. In the following, given a contig *c* ∈ *C* we denote by *F*_*c*_ the set of fragments into which *c* has been partitioned, while, given a fragment *f* ∈ *F*, we denote by *C*_*f*_ the set of contigs of which *f* is a part. Note that if *f* is shared then |*C*_*f*_ | *>* 1, otherwise (i.e., if *f* is private) |*C*_*f*_ | = 1. Please refer to Figure 1.1 and 1.2 for a visual representation of these concepts.

**Figure 1:**
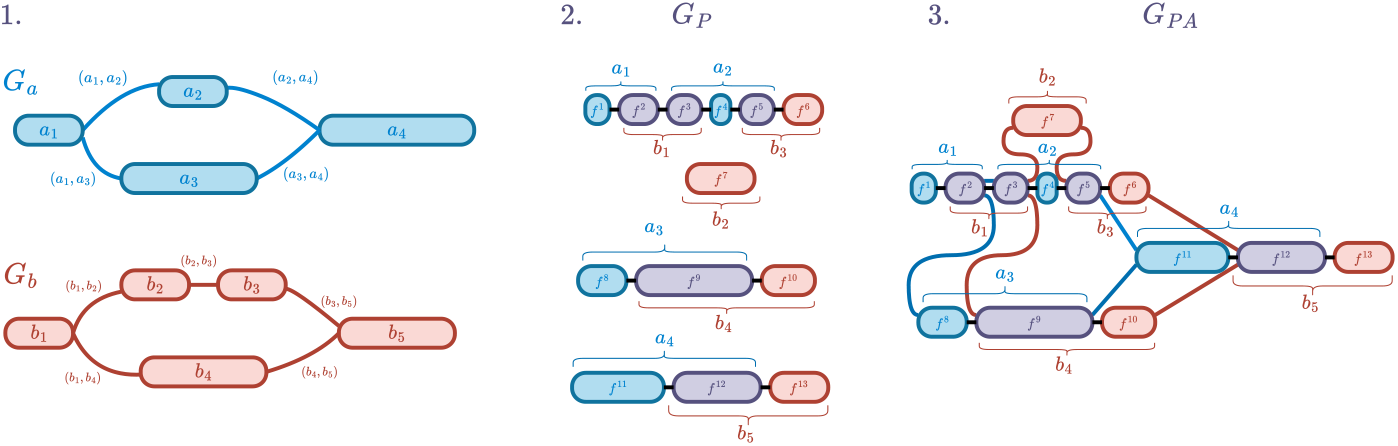
An illustration of how `Pangebin` builds the pangenome-assembly graph. 1. Input assembly graphs *G*_*a*_ = (*C*_*a*_, *E*_*a*_) and *G*_*b*_ = (*C*_*b*_, *E*_*b*_). 2. The pangenome graph *G*_*p*_ = (*F, E*_*p*_) obtained by overlapping the contigs in *C* = *C*_*a*_ ∪ *C*_*b*_. Original contigs in *C* have been fragmented into smaller nodes (for example, contig *a*_1_ has been fragmented into 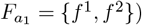. Overlaps are shown by shared fragments colored in violet, while private fragments are colored in blue (red, respectively) if obtained from *G*_*a*_ (*G*_*b*_, respectively). 3. The pangenome-assembly graph *G*_*PA*_ = (*F, E*_*asm*_, *E*_*p*_) obtained by adding the edges of the input assembly graphs. Note, for example, that edge (*a*_1_, *a*_3_) of *G*_*a*_ induces edge 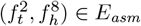, linking the tail of the last fragment of *a*_1_ with the head of the first fragment of *a*_3_.

### 2.2 The pangenome-assembly graph *G*_*PA*_: from graph *G*_*p*_ to a consensus of assembly graphs

The pangenome graph *G*_*p*_ contains only the information about fragments and how they are contiguous within the given contigs. This step transforms the pangenome graph *G*_*p*_ into the pangenome-assembly graph *G*_*PA*_ by incorporating the edges *e* ∈ *E*_*a*_ ∪ *E*_*b*_ of the input assembly graphs *G*_*a*_ = (*C*_*a*_, *E*_*a*_) and *G*_*b*_ = (*C*_*b*_, *E*_*b*_). Moreover, for each original contig *c* ∈ *C*_*a*_ ∪ *C*_*b*_ additional metadata present in the assembly graphs, such as coverage and GC-content, are added to the pangenome-assembly graph as detailed below. Formally, the pangenome-assembly graph *G*_*PA*_ = (*F, E*_*asm*_, *E*_*p*_) is obtained by augmenting the pangenome graph *G*_*p*_ = (*F, E*_*p*_) with the following elements:

- *assembly edges*: edges in *E*_*asm*_ linking together fragments of different contigs from the same assembly graph;
- *node features*: each fragment *f* ∈ *F* is annotated with features such as length (*l*_*f*_), GC-content (*gc*_*f*_), and coverage (*rc*_*f*_).

We first detail the addition of assembly edges. For each edge 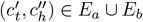, we add the edge 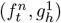 to *E*_*asm*_, where *f*^*n*^ is the last fragment of contig *c*^*′*^, while *g*^1^ is the first fragment of contig *c*^*′′*^ in graph *G*_*p*_. Figure 1.3 provides an example of the final pangenome assembly graph *G*_*PA*_ obtained by simply adding to *G*_*p*_ in Figure 1.2 each edge *e* in the set *E*_*asm*_, from *G*_*a*_ and *G*_*b*_ in Figure 1.1

Node features are then added to *G*_*PA*_ as follows. For each contig *c* ∈ *C*, we transfer its features to the corresponding fragments *f* ∈ *F*_*c*_ ⊆ *F*. Note that some fragments are shared by multiple contigs, so we compute the features of each fragment *f* as the mean of the features of the contigs in *C*_*f*_ that are sharing a particular fragment *f*. Specifically, we compute the coverage *rc*_*f*_ (GC-content *gc*_*f*_, respectively) as the mean of the coverage (GC-content, respectively) of the contigs that overlap on that particular fragment *f* (i.e., contigs of set *C*_*f*_).

### 2.3 PlasBin-flow on Pangenome-assembly graphs

`PlasBin-flow` is a binning tool that works iteratively by solving a Mixed-Integer Linear Programming problem. Here we describe how to have a version of `PlasBin-flow` that solves the binning problem on pangenome-assembly graphs. As already observed, since `PlasBin-flow` solves a Mixed-Integer Linear Programming problem, it can be naturally modified to include constraints related to the fact that the pangenome-assembly graph combines fragments coming from different assembly graphs. In particular, we show how to take into account that some fragments are not supported by both assemblers by introducing a penalty score that contributes to the final objective function of the MILP formulation. As detailed before, the features of nodes already take into account that a fragment inherits the plasmidness, coverage, and GC-content from the contigs that share it. All these score values are an essential part of the MILP.

The Mixed-Integer Linear Programming for the pangenome works by identifying a set of fragments in the graph that could belong to the same plasmid of a particular bacterium, calling that a *plasmid bin*. Fragments belonging to this set are identified starting from a *seed fragment* : we chose seed fragments as nodes in the graphs being at least 1000bp long and with a gene density score of at least0.5. The choice of these values is based on lowering the average values used in `PlasBin-flow` on a single assembly graph, since we take into account that fragments come from fragmentation of contigs. Subsequently, fragments from the plasmid bin are removed from the graph, repeating the process until no more seed fragments are left in the graph. On the pangenome-assembly graph, a plasmid bin can be defined as a connected subgraph that contains at least one seed fragment.

Each plasmid bin is then computed identifying a subgraph through the optimization problem defined as a linear combination of (1) a flow value ℱ, that accounts for sequencing coverage of its fragments, (2) a penalty value *GC*, that accounts for the probability of each fragment’s gene content to diverge from the overall GC-content of the plasmid bin, (3) a gene density penalty *GD*, that accounts for the probability of a fragment of being a plasmid, given a database of plasmid genes, and (4) a private penalty *PP*, that is a score we added in our adapted version of `PlasBin-flow` for penalizing *private* fragments.

The adapted MILP is hence defined as a maximization of ℱ + *GC* + *GD* + *PP*. The main decision variable is *x*_*f*_, representing the whole fragment *f*. Then we have a continuous variable *f* (*e*), which encodes for the flow through each edge *e* (accounting for coverage) and the flow ℱ that describes the overall flow value in the graph *G*_*PA*_. Each plasmid is then identified by a sub-graph *G*_*PA*_(*p*) that maximizes the objective function by the above constraints.

Formally, we have:

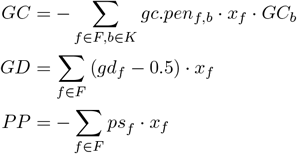

In detail, *gc*.*pen*_*f,b*_ indicates the probability of the fragment *f* to deviate from the GC-content interval of the current plasmid bin (*GC*_*b*_), where intervals are defined by the set *K* = *{*[0, 0.4], [0.4, 0.45],

[0.45, 0.5], [0.5, 0.55], [0.55, 0.6], [0.6, 1]*}*. Moreover, *b* is a binary decision variable that describes the chosen *k*-interval in *K. gd*_*f*_ indicates the probability, from 0 to 1, that fragment *f* is a true plasmid, and it is added in the objective function by subtracting 0.5, hence penalizing non-plasmid fragments and supporting plasmid fragments. The last term is the private score *ps*_*f*_ that we added as a penalty term with the final aim of including fewer private fragments in the solution. In fact, since our approach is based on the idea of building and analyzing a consensus graph, we want to penalize fragments supported by only one assembler since those may represent errors during the assembly process. Moreover, the higher the similarity is between two matched contigs, shorter are the unmatched sequences, i.e. the private fragments produced. In order to account also for this fact, we wanted to penalize less shorter fragments, i.e. shorter assembly errors, and penalize more larger private fragments, up to 1000bp.

The private score *ps*_*f*_ is hence defined as follows:

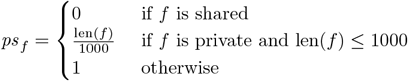

## 3 Evaluation

We design an experimental analysis with a twofold objective: first, we aim to assess the impact of a specific assembler on precision and recall of a plasmid binning tool, and second, we evaluate the performance of our tool `Pangebin` to mitigate the negative impact of a given assembler on a sample, measuring the accuracy of predictions over the pangenome-assembly graph. We run experiments over a dataset of 48 short-read bacterial isolates, 40 *Klebsiella pneumoniae* and 8 *Escherichia coli*. The 48 samples have been extracted out of 112 samples of the test set of Plasgraph 2 [18] (originally sampled from the Robertson and Nash dataset [4]) since they were the only samples fully annotated with plasmid information in the RefSeq NCBI [19] database. Indeed, only on these samples we could produce the ground-truth based on a previous annotation. The samples have been assembled using Skesa [14] and Unicycler [13], obtaining a dataset of 48 pairs of assembly graphs. The evaluation is done by computing the ground truth based on the assembly graphs on which the specific tool was run to get the plasmid bins. In other words, a plasmid binning tool with input a given assembly graph is evaluated by first assigning to nodes (i.e., contigs) of the graph the *labels* (i.e., plasmid or chromosomal) and *identifiers* (i.e., the specific plasmid) based on the reference database. More precisely, the sequence of each contig of these assemblies is matched against the reference using `BLAST` [20], assigning them a `plasmid` (`chromosome`) label if the sequence was matching with at least 95% identity a `plasmid` (`chromosome`) sequence, preferring the best match in case of multiple matches, or `unlabeled` if no match was found. Moreover, based on the specific plasmid annotation given in the reference, we label each node with a plasmid identifier to distinguish them. Note that unlabeled nodes are removed in the evaluation step.

To compute the ground truth on the pangenome-assembly graph, we mapped the ground truth of each assembly graph to the nodes in the pangenome, labeling private fragments with the same label as the original contig they came from, and shared fragment *f* with the label assigned to the longest contig in *C*_*f*_. In the end, we have a pair of ground truth labels for each node in the pangenome-assembly graph, each one coming from the Skesa and Unicycler ground truth respectively.

We compared the performance of `Pangebin` on the proposed dataset against the original `PlasBin-flow` and against `gplas`. `PlasBin-flow` and `gplas` have been separately applied to both Unicycler and Skesa assembly graphs. The comparison has been done on labeling and binning performances, where *labeling* measures the ability of the tool to correctly identify plasmid nodes and *binning* measures the accuracy in outputting the plasmid bins, i.e., clusters of nodes that were recognized to belong to the same original plasmid. Given a set of nodes *N*, we define ‖*N* ‖ to be the total length of sequences assigned to each node in *N*. Precision, recall, and F1-score for labeling have been evaluated considering as solution the superset of nodes *S* obtained by the union of each output bin, weighting the accuracy scores by ‖*S* ‖. More precisely, we consider:

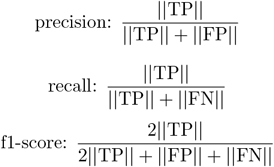

where TP (true positive) represents the set of nodes correctly identified as plasmids, FP (false positive) represents the set of nodes wrongly identified as plasmids, and FN (false negatives) is the set of nodes labeled as plasmid in the ground truth that are missing in the set of nodes in solution *S*. Note that we are considering each accuracy metric weighted by the size of the nodes considered in each set TP, FP and FN. Binning has been evaluated using the accuracy metrics provided by PlasEval [21]. PlasEval evaluates the performance mapping nodes in each bins to sets of nodes in the ground truth that share the same plasmid identifier: each distinct plasmid sequence *p* in the reference corresponds to a set *R*_*p*_ of nodes of the assembly graph that match to *p*. Then, for each predicted bin *b* (output solution of a given tool), *P*_*b*_ is defined to be the set of nodes that the tool assigns to plasmid *b*. In order to compute binning precision, we need to assign to each predicted bin *b* one reference plasmid *p*, i.e., the one that maximizes |*P*_*b*_ ∩ *R*_*p*_|. Then, the precision for the bin *b* is computed as the ratio of |*P*_*b*_ ∩ *R*_*p*_|over |*P*_*b*_|. The overall binning precision of the sample is computed as the mean of the precision of each bin *b*, weighted by their size |*b*|. Similarly, the binning recall is computed assigning to each reference plasmid *p* a predicted bin *b* such that the |*P*_*b*_ ∩ *R*_*p*_| is maximized. The recall of the plasmid *p* is then computed as the ratio of |*P*_*b*_ ∩ *R*_*p*_| over |*R*_*p*_|. F1-Score of the binning is computed as the harmonic mean of precision and recall, weighted by the overall length of the bins.

First we evaluated how `gplas` and `PlasBin-flow` are performing in terms of accuracy over a given assembly graph. In Table 1 and Table 2 we reported, respectively, the labeling and binning mean accuracy scores observed for the two tools run on each pair of input assembly graphs. Both `PlasBin-flow` and `gplas` have been run with default parameters, and the latter in particular was ran using `mlplasmids` as preliminary contig classification tool. We observed that overall, given a plasmid binning tool, it is not completely clear which assembler will perform better over a given sample and this is particularly true for `PlasBin-flow` that has higher average precision (+0.27) on Skesa assemblies and slightly higher average recall (+0.04) using Unicycler assemblies. A similar situation is observed for binning accuracy, where `PlasBin-flow` performs better in precision with Skesa assemblies (+0.22) and better in recall with Unicycler assemblies (+0.12). `gplas`, instead, performs better when used along with Unicycler assembly graphs, both for labeling and binning. Note that `gplas` was not originally intended to work with assembly graphs different from Unicycler, and in order to have this evaluation, we adapted it to handle Skesa assembly graphs. The distribution of precision, recall, and F1-score is presented in Figure 2 and Figure 3 for labeling and binning, respectively.

**Figure 2:**
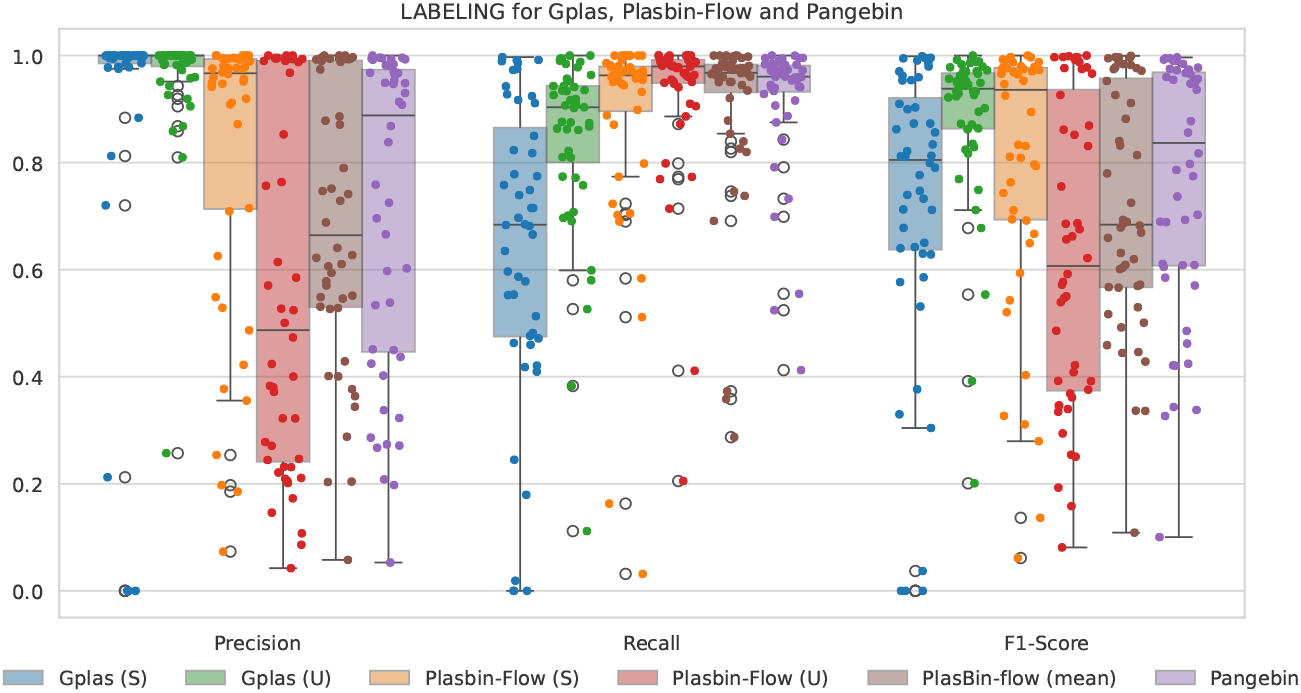
Labeling accuracy of `Pangebin` against `gplas` and `PlasBin-flow` run on Unicycler (U) and Skesa (S) assembly, and against the mean scores of `PlasBin-flow` run on both assemblies.

**Figure 3:**
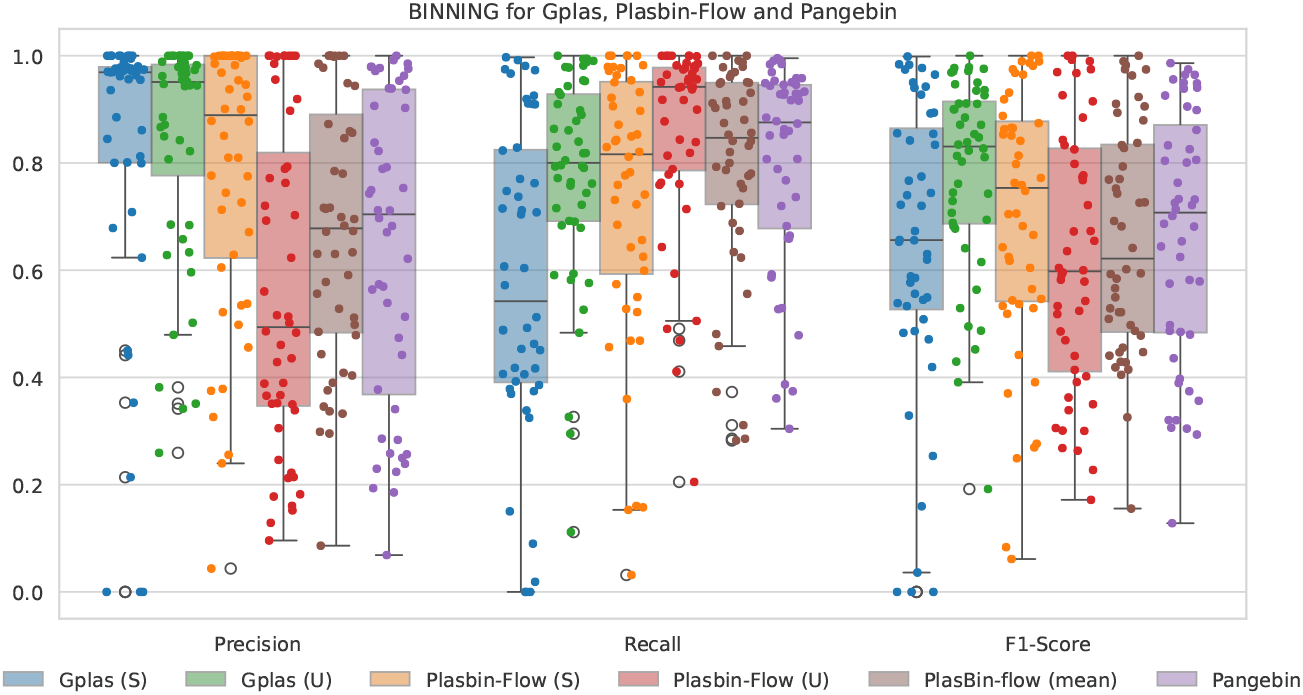
Binning accuracy of `Pangebin` against `gplas` and `PlasBin-flow` run on Unicycler and Skesa assembly, and against the mean scores of `PlasBin-flow` run on both assemblies.

**Table 1:** Labeling performances for `Pangebin`, `PlasBin-flow`, and `gplas` (mean *±* std. dev.). The best overall performances are underlined and the best performances between `Pangebin` and `PlasBin-flow` mean performances are in bold.

| Tool | Assembler | Precision | Recall | F1-Score |
| --- | --- | --- | --- | --- |
| Pangebin | “pangenome” | **0.718 ± 0.297** | **0.917 ± 0.127** | **0.764 ± 0.238** |
| PlasBin-flow | “mean” | 0.690 ± 0.265 | 0.906 ± 0.165 | 0.712 ± 0.224 |
| PlasBin-flow | Unicycler | 0.558 ± 0.341 | 0.927 ± 0.148 | 0.628 ± 0.283 |
| PlasBin-flow | Skesa | 0.823 ± 0.269 | 0.884 ± 0.202 | 0.797 ± 0.250 |
| gplas | Unicycler | <u>0.963 ± 0.112</u> | 0.843 ± 0.172 | <u>0.882 ± 0.155</u> |
| gplas | Skesa | 0.906 ± 0.266 | 0.640 ± 0.283 | 0.721 ± 0.278 |

**Table 2:**
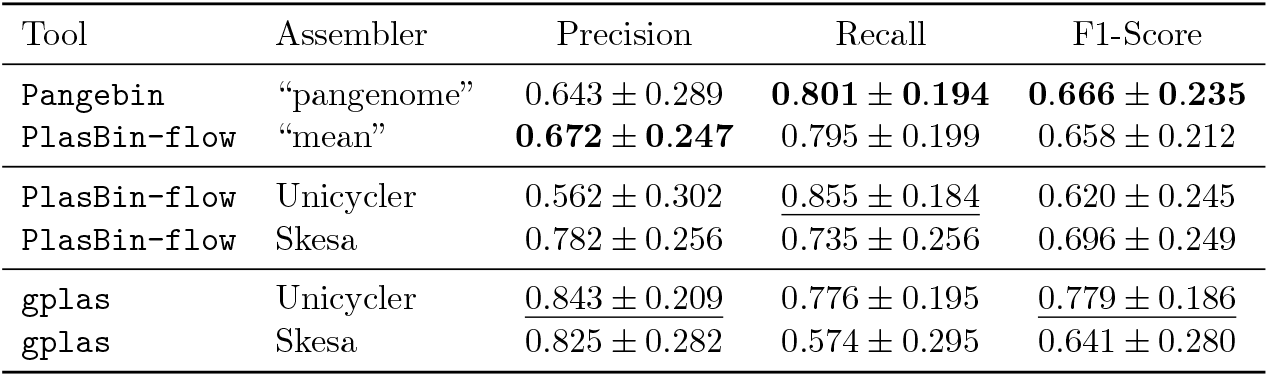
Binning performances for `Pangebin`, `PlasBin-flow`, and `gplas` (mean *±* std. dev.). The best overall performances are underlined and the best performances between `Pangebin` and `PlasBin-flow` mean performances are in bold.

Second, we evaluated `Pangebin` labeling and binning accuracy compared to `PlasBin-flow`, observing that the former performs better in labeling contigs if compared to the mean accuracy scores obtained on the pair of assembly graphs of each given sample, scoring +0.018 in precision, +0.011 in recall, and +0.052 in F1; it also performs slightly better in binning recall (+0.06) and F1-Score (+0.08). Note also that `Pangebin` obtains the second-best score in labeling and binning recall (0.917 and 0.801 respectively). Moreover, it is true that the mean-assembly scores are not directly obtained by evaluating the output of `PlasBin-flow` run over a hypothetical mean-assembly that does not in fact exist, but those scores are instead a mere computed mean of two evaluations. The pangenome-assembly graph, in this sense, can be thought as a mean-assembly graph that aims at performing better than the original pair of assembly graphs on average. Note that `Pangebin` is not able to reach the best scores obtained by a specific assembler over a particular metric, but on the other hand `Pangebin` always improves the worst solution: `Pangebin` gains +0.081 in binning precision compared to `PlasBin-flow` over Unicycler assemblies and gains +0.066 in binning recall compared to `PlasBin-flow` over Skesa assemblies.

These results justify the use of the pangenome-assembly graph to smooth the effect of the assembly graph over the performance of a given tool. To further illustrate this, in Figure 4 we compared binning precision and recall achieved by `Pangebin` and `PlasBin-flow` (run on Skesa and Unicycler assemblies) on each sample. We also compared the mean performances over the two types of assembly graph. The last row of plots shows indeed that `Pangebin` provides a more balanced and robust solution, in line with the mean performances of `PlasBin-flow`. In the end, this evaluation shows that the pangenome-assembly graph can be a good way of mitigate the effect of discordant assembly graphs over the binning performance of a specific plasmid binning tool.

**Figure 4:**
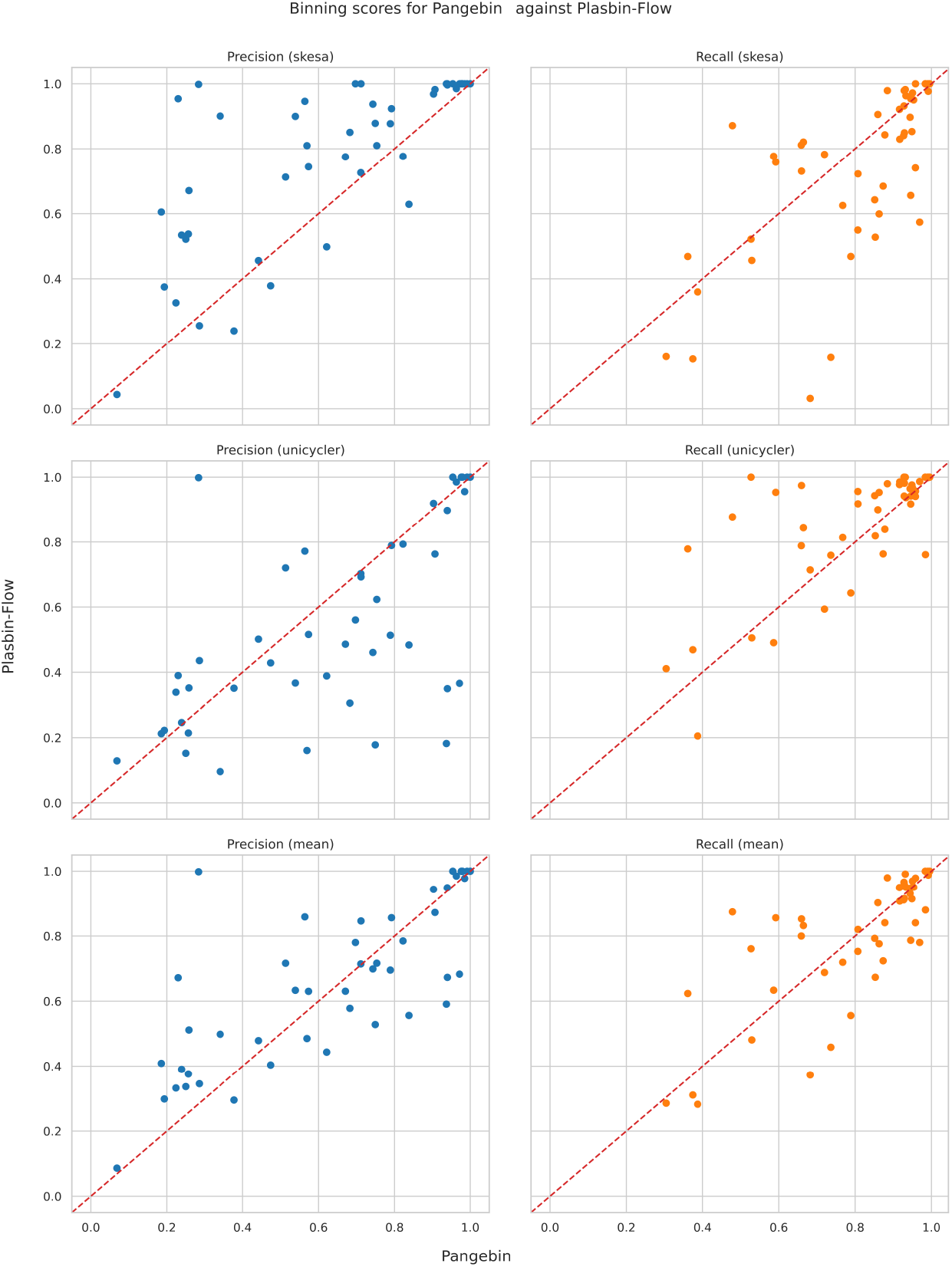
Scatterplot of binning precision and recall of `Pangebin` against `PlasBin-flow`, run on Skesa and Unicycler; mean performances are reported in the bottom plots. Dots below the red diagonal represent samples performing better with `Pangebin`

## 4 Conclusions

In this paper, our main contribution is the proposal of a pangenome-assembly approach for plasmid binning that aims at lessening the impact of assembly errors on the accuracy of plasmid binning tools. The main idea implemented in our tool `Pangebin` is combining the assembly from different tools from the same short-read data by leveraging the recent advances in pangenome analysis. Indeed, without a priori knowledge of the best assembly, a pangenome-assembly approach is expected to find a consensus between the results achieved by a plasmid binning tool over the different graphs. As summarized below, our paper demonstrates that our approach optimizes accuracy in plasmid binning across different assembly graphs. An interesting future research direction opened by this work is exploring the pangenome-assembly approach for other tasks on assembly graphs derived from the same short-read data.

`Pangebin` integrates a modified version of `PlasBin-flow` that has been adapted to work on pangenome-assembly graphs with overall good results. In our experimental analysis, we show that, without prior knowledge of the performance of a prediction tool over a given assembly graph, `Pangebin` is able to obtain results that are close to the best performance achievable on one of the assembly graphs, in both labeling and binning metrics. We compared our tool against `PlasBin-flow` showing that, on average, our solution performs slightly better, with the advantage that we always get a good solution on average. By using `Pangebin` we are able to improve over the precision of `PlasBin-flow` achieved with Unicycler, while also improving the recall achieved with Skesa.

Furthermore, we presented a comparison between `PlasBin-flow` and `gplas` to support the fact that different combinations of binning tool and assembly graphs produce indeed very diverse results for a given sample. We observed that `PlasBin-flow` tends to have a higher recall at the expense of lower precision when used in combination with Unicycler, while we observe the opposite behavior when used with Skesa. `gplas`, on the other hand, performs better when used along with Unicycler assemblies, but shows lower recall in general compared to `PlasBin-flow`. Moreover, `gplas` cannot be easily adapted to work with pangenome-assembly graphs, since it relies on specific graph topology to compute plasmid bins. More in detail, `gplas` computes plasmid bins starting from seed contigs (nodes) that have in- and out-degree both equal to 1. As is clear from Figure 1, every shared contig in the pangenome-assembly graph has at least in- or out-degree greater than 1 and thus cannot be considered a seed contig. In fact, despite we were able to adapt `gplas` to pangenome-assembly graphs, most samples produced no results, caused by the lack of seed contigs in the pangenome-assembly. For this reason, we decided not to present such results in this manuscript, since those would not add any value to the comparison. Given that, further investigation on how to adapt `gplas` to the different topology of the pangenome-assembly graph is an interesting future development.

The key ingredient that allows `Pangebin` to achieve such results is indeed the fragmentation of contigs into shared fragments, that provides a comprehensive view over the sample. Nevertheless, in our experimental analysis, we also observed that the fragmentation comes with the side effect of splitting a plasmid into multiple bins; this is also true for the original formulation of `PlasBin-flow` and, by how the binning evaluation is designed, this fact indeed penalizes the recall metric. In the end, we expect to further improve the accuracy of `Pangebin` over `PlasBin-flow` by modeling in the MILP the specific fragmentation coming from contigs that represent the same plasmid by adding proper corrective values to the objective functions.

## 6 Summary

- Assembly graphs produced by different tools on the same bacterial sample may vary significantly, creating challenges for plasmid binning.
- Pangebin is a new method that leverages pangenome graphs to reveal similarities between contigs from different assemblies to guide plasmid binning across multiple assembly graphs from the same short-read sample.
- Pangebin, without prior knowledge of the best assembly, can identify a consensus among the results of Plasbin-Flow binning tool applied to different graphs.
- The Pangenome-assembly approach to mix together two -or potentially more-assembly graphs has the potentiality to guide downstream analysis in absence of a priori knowledge of the best assembly.

